# One week of chronic sleep debt does not affect decision-making processes in a mouse version of the Iowa Gambling Task

**DOI:** 10.1101/691246

**Authors:** Elsa Pittaras, Sylvie Granon, Arnaud Rabat

**Affiliations:** Unité Fatigue et Vigilance, Institut de Recherche Biomédicale des armées, 91223, Brétigny-sur-Orge cedex. Equipe d’accueil VIgilance FAtigue et SOMmeil (VIFASOM) EA 7330 – Université Paris 5 Descartes, 75005, Paris, France; Equipe ‘Neurobiologie de la prise de décision’, Neuro-PSI, CNRS UMR 9197, 91400, Orsay, France; Stanford University, United-States

## Abstract

Socio-professional pressures push people to sleep less which leads to chronic sleep debt (CSD) for a significant percentage of the population. Although the health consequences of CSD are well known, research shows that high-level cognitive processes in humans are more affected by acute sleep debt (ASD) rather than CSD (Drake et al., 2001). We have previously shown that ASD has deleterious effects on decision-making in mice and that some mice were more sensitive to ASD than others (Pittaras et al., 2018) by using a rodent version of the Iowa Gambling Task (Bechara et al., 1994). In this study, we showed that, as in humans, CSD has fewer effects on decision-making compared to ASD. We hypothesize that this observation was due to the set-up of a compensatory mechanism.

## Introduction

Socio-professional pressures often lead people to decrease the amount of time that they spent in bed. Indeed, a recent survey from the National Sleep Foundation showed that sleep is not a priority among other personal activities: only 10% of people picked sleep as the most important items compared to 35% who chose physical fitness, 27% who selected work, 17% who preferred hobbies and 9% who picked their social life (Survey 2018 from National Sleep Foundation). These choices have led to a decreased amount of sleep and chronic sleep debt (CSD) for a large percentage of the population. Sleep loss refers to sleep of a shorter duration than the average basal need of 7 to 8 hours per night. The health consequences of sleep loss are far from benign. Indeed, the cumulative long-term effects of sleep loss have been associated with a wide range of deleterious health consequences including heart attack or stroke but also with cognitive deficits (Khan et al., 2017; Palma et al., 2013; Killgore et al., 2009). It has been shown that sleep loss leads to excessive daytime sleepiness, depressed mood and poor memory or concentration (Dinges et al., 1997). However, studies looking at cognitive effects are less abundant and only a few studies have focused on high-level cognitive processes that require a large brain network such as decision-making (Khazaie et al., 2010; Killgore et al., 2006).

We previously showed that acute sleep debt leads to decision-making deficits in mice by inducing less flexibility and that ASD affects more extreme decision-making profiles (Pittaras et al., 2018). We also showed that dopamine and serotonin modifications in the caudate putamen and the orbitofrontal cortex could explain the decision-making deficits and the lack of flexibility (Pittaras et al., 2018). It has been shown in humans that memory deficits are more significant after one night of acute sleep debt than after two nights of chronic sleep debt (Drake et al., 2001) and that less than 6 hours of sleep over 5 days’ period has no effect on decision making during the Iowa Gambling Task (Khazaie et al., 2010) but an acute sleep deprivation altered decision making during this task (Killgore et al., 2006). Therefore, we asked the following question: is a chronic sleep debt lasting one week less deleterious than 23 hours of sleep loss in mice regarding decision-making deficits as it was observed in humans?

To answer this question, we used a decision-making task under ambiguity that we developed in mice (Pittaras et al., 2016) and completed this task under chronic sleep debt. We showed that, as in humans, an acute sleep debt is more deleterious than a chronic sleep debt during decision-making processes.

## Materials and methods

### Animals

49 mice were used for the Mouse Gambling Task (MGT). The mice were bred in Charles’ River facilities (Orleans, France) and were 3 months old at the beginning of the experiments. They were housed in a collective cage of 3 or 4 in a temperature-controlled room (22 ± 2 °C) with a fixed light/dark cycle (light on at 8:00 a.m. and light off at 8:00 p.m.). All behavioral experiments were performed during the light cycle between 9:00 a.m. and 5:30 p.m. For the Mouse Gambling Task, the mice were food deprived (maintenance at 85 % of the free feeding weight) but always received water *ad libitum*.

### Mouse Gambling Task (detailed protocol in Pittaras et al., 2019)

#### Habituation in operant chambers

The aim was to habituate the mice to the experimenter, to allow the mice to eat pellets in an area other than their home cage, to train them to perform a behavioral action in order to receive food pellets and to stabilize mice weight at 85% of their free-feeding weight by the end of this habituation in operant chambers.

Each mouse did 1 session per day for 10 days. We changed the running order of mice each day to avoid any time effect. One session lasted 30 minutes. During these sessions, the central hole was the only hole available in the operant chamber which contained 5 holes and a magazine. When a mouse did a nose poke in the central hole, one food pellet was distributed. The mouse then had to visit the magazine to eat the pellet and a new trial began.

Results of this habituation are not shown here. We observed an increase of pellets consumption across days for all mice.

#### MGT

The task took place in a maze with four transparent arms (20 cm long × 10 cm wide) containing an opaque start box (20 cm × 20 cm) and a choice area (**Figure 1A**). We used standard food pellets as a reward (dustless Precision Pellets, Grain-based, 20 mg, BioServ® -New-Jersey) and food pellets previously steeped in a 180 mM solution of quinine as penalty (VandenBos et al. 2006). The quinine pellets were unpalatable but not unedible. Each mouse performed 10 trials in the morning and 10 trials in the afternoon for 5 days (1 day = 20 trials = 1 session; 5 days = 5 sessions = 100 trials as with the human task; Bechara et al. 1994). There were 4 different arms: 2 that gave access to long-term “*advantageous*” choices and 2 others that gave access to long-term “*disadvantageous”* choices. In the long-term “*advantageous*” arms, mice could find 1 pellet (“small reward”, as the $50 in the IGT) before a bottle cap containing 3 or 4: food pellets on 18 trials out of 20 and quinine pellets for 2 trials out of 20. In the “*disadvantageous”* arms mice could find 2 food pellets (“large reward”, as the $100 in the IGT) before a bottle cap containing 4 or 5: quinine pellets 19 trials out of 20 and food pellets on 1 trial out of 20 (**Figure 1A**). “*Advantageous*” choices are at first less attractive because of their small immediate reward (1 pellet) whereas “*disadvantageous*” choices are at first more attractive because of their access to large immediate reward (2 pellets). Despite the immediate less attractive reward, “*advantageous*” choices are more advantageous in the long term because animals more often found food pellets than quinine pellets. Likewise, “*disadvantageous*” choices are less advantageous in the long term because animals more often found quinine pellets than food pellets (**Figure 1A**). Thus, mice had to favor a small immediate reward (“*advantageous”* choices) to obtain as many pellets as possible by the end of the day.

**Figure 1:**
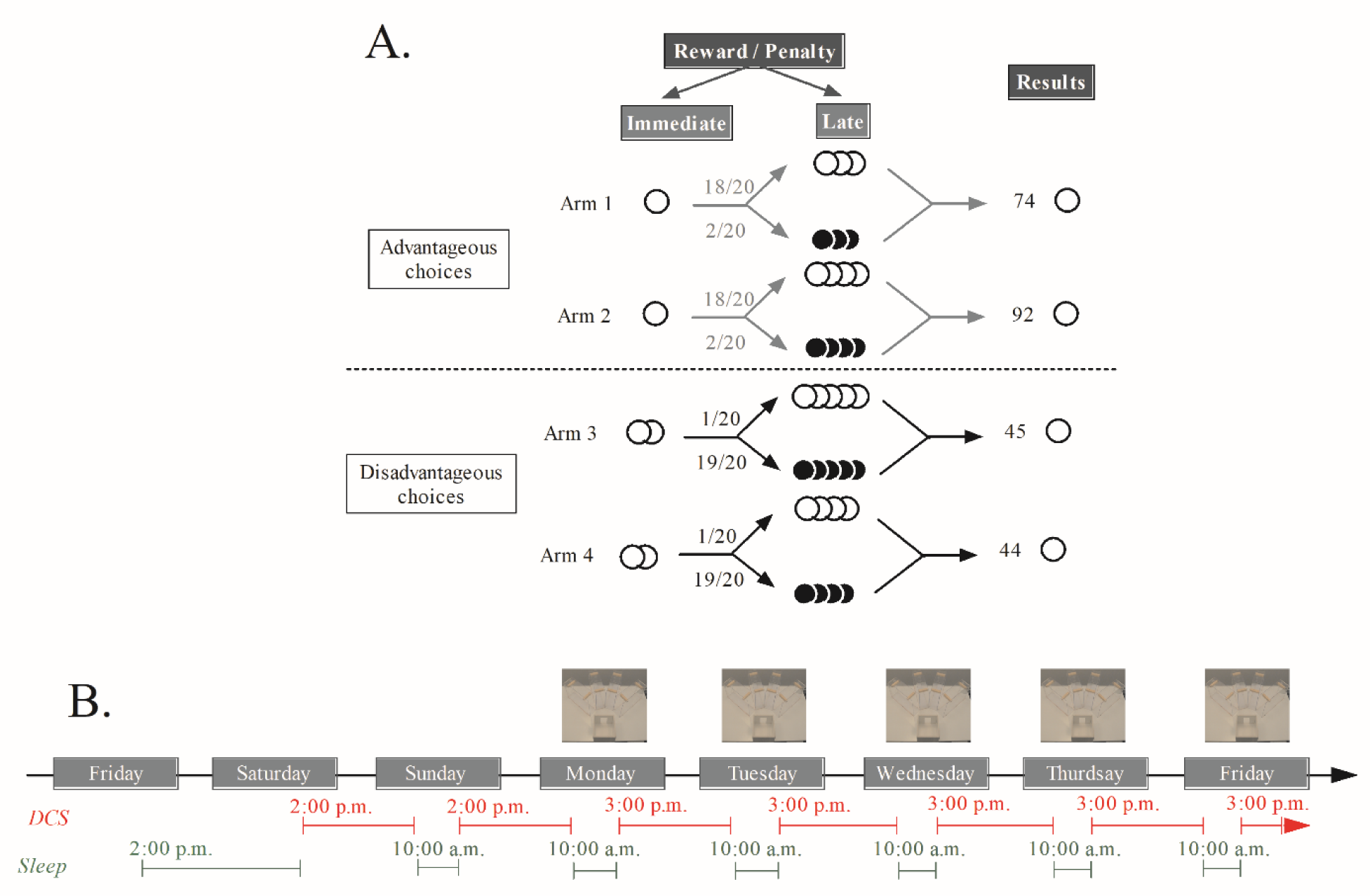
**A. Schematic representation of the MGT experimental design.** White circles represent food pellets and black circles represent quinine pellets. Advantageous choices gave access to one food pellet and disadvantageous choices gave access to two food pellets. Then the mouse can find three or four food pellets (18/20) or quinine pellets (2/20) in advantageous choices and four or five food pellets (1/20) or quinine pellets (19/20) in disadvantageous choices. We distinguished advantageous choices from disadvantageous ones because mice earned more pellets (74 or 92 pellets vs. 45 or 44 pellets) after 20 trials by choosing the advantageous ones. **B. Timeline of the Chronic Sleep Debt (CSD).** Mice were placed in the cylinder at the end of the first Friday for habituation for 24 hours. The CSD started on Saturday at 2:00 pm and lasted for 6 days. During the day without MGT, the sleep deprivation began at 2:00 pm and stopped at 10:00 am. Mice were thus able to sleep for 4 hours. During days with MGT, the cylinders were activated at 3:00 pm and stopped at 10:00 am. Mice were thus able to sleep for one more hour as the completion of the MGT might bother them.

Between each trial, the maze was cleaned with distilled water and, between each mouse, it was cleaned with a 10% alcohol solution. During the first session, animals were put into the maze for 5 minutes with food pellets scattered everywhere (habituation in the maze). If a mouse did not eat any food pellets during the first habituation, a second 5-minute habituation period was conducted. For the following sessions, habituation lasted only 2 minutes without food pellets in the maze. At the beginning of each trial, the mouse was placed in an opaque tube in the starting box to avoid directing the future choice of the animal. After around 5 sec., we removed the opaque tube and let the animal freely choose one arm of the maze. We measured the time the mouse took to choose one arm (i.e. when the animal crossed 1/3 of the arm) and we scored the chosen arm and the amount of pellets consumed (pellets earned). A rigidity score was calculated by measuring how many time the animal chose the same arm. For example, the rigidity score was 25% if animals chose as many advantageous options as disadvantageous ones. A 50% score reflected that the animal chose one arm twice more than the others and a 75% score indicated that the animal chose ¾ of the time the same arm than the other.

MGT is organized in two phases: a *phase of exploration* (discovery of options) and a *phase of exploitation* (better knowledge of options). Usually, during *the exploration phase (day 1 and 2)*, mice do not show any preference between options. However, during the *exploitation phase (day 3, 4 and 5)* preferences for long-term advantageous options emerge with inter-individuals’ differences i.e. some animals show a marked preference for long-term advantageous options (safe) or don’t show any preference for advantageous options (risky) while others make choices in between (average).

### Chronic sleep debt

Animals were sleep deprived using an apparatus (Chauveau et al. 2014) with a transparent cylinder (PVC, 45 cm height) on a shaking platform (diameter 30 cm) (Viewpoint®). Animals stayed awake because the platform bounced at random frequency (stimulation every 0.15 to 0.20 min), intensity (1cm height) and duration (30 to 45 ms). Moreover, the number (3 to 6) and the duration of stimulation (0 to 100 ms of intervals) were randomized to ensure that the mice did not acclimate. Software controlled the shaking parameters (Viewpoint, France) during the experiment. We put 4 mice per cylinder. They were able to drink and eat.

Mice were put in the apparatus for 7 days: 1 day of habitutation (start at 2:00 p.m. on Friday) and 6 days of chronic sleep debt. Control mice were also put in the apparatus during the 7 days but we never activated the apparatus during that period. Chronic sleep debt began on Saturday at 2:00 p.m. and the MGT began on Monday morning. Mice could sleep between 10:00 a.m. and 2:00 p.m. on Sunday and between 10:00 a.m. and 3:00 p.m. on all others days. Thus, CSD lasted 20 hours on Saturday and 19 hours on all the other nights. Indeed, as the MGT was done during the morning and the afternoon, all animals were often bothered upon completion of the MGT. Therefore, we choose to let the mice sleep for 1 more hours during the Mouse Gambling Task (**Figure 1B**).

### Statistical analysis

Statistical analyses were performed using Statview software. For the data that showed normal distribution (Shapiro-Wilkonson test) and passed equal variance tests (F test), statistical analyses were performed using t-test. When data did not show normal distribution or pass equal variance tests, statistical analyses were performed using ranked signs Wilcoxon task, Kruskal Wallis or Mann-Whitney U tests. Bonferroni correction was applied for multiple comparisons. Results were reported as means ± SEM. P values ≤ 0.05 were considered statistically significant except when Bonferroni correction was necessary.

#### Sub-groups formation

To distribute animals among groups according to their MGT preference, we calculated, for each mouse, the mean of the percentage of advantageous choices for the 30 last trials, i.e. when preference was stable, and used a k-mean clustering analysis with Statistica® software (version 12, Pittaras et al., 2016, Timmerman et al., 2013). With this objective method, each animal belonged to a set that had the closest mean to its own preference value. Animals were separated into three subgroups: animals that chose a majority of advantageous options at the end of the experiment, called “*safe*”; animals that explored options until the end of the experiment, called “*risky*”; and animals that maintained some exploration of available options but favored advantageous options, called “*average*” (Pittaras et al., 2016).

## Results

### Global effects of CSD

As we observed previously (Pittaras et al., 2016), control mice preferentially chose long term advantageous options at the end of the MGT (day 1 vs. day 5: t=−2.368, p=0.0281). Interestingly, we observed the same progression of the percentage of advantageous choices for CSD mice (day 1 vs. day 5: t=−2.518, p=0.0158). Indeed, CSD did not affect mice preference: control mice and CSD mice never differed from each other (day 1: t=−0.140, p=0.8897; day 2: t=−0.583, p=0.5625; day 3: U=341.500, p=0.3372; day 4: U=220.000, p=0.1349; day 5: t=0.002, p=0.9982).

Control animals as well as CSD mice chose long term advantageous options faster as the days of MGT progressed (Control mice: day 1 *vs.* day 5: Z = −4.015, p < 0.0001; CSD mice: day 1 *vs.* day 5: Z = −4.031; p < 0.0001). Moreover, CSD mice were faster than control mice during all days of MGT expect the fourth day (day 1: U=138.000, p=0.0016; day 2: t=2.810, p=0.0072; day 3: U=152.000, p=0.0041; day 4: U=216.000, p=0.1151; day 5: U=168.000, p=0.0109).

Interestingly, despite the lack of effect of CSD on mice’s preferences during the MGT, we observed an effect of CSD on mice rigidity scores during this task. Indeed, only CSD mice had a higher score of rigidity at the end of the MGT compared to the beginning of this task (CSD mice: 2 first *vs.* 2 last days: t=−2.277, p = 0.0309; Control mice: 2 first *vs.* 2 last days: t=−1.724, p=0.1001). However, no difference between CSD and control mice was observed (2 first: t=−0.807, p=0.4240; 2 last days: t=−0.993, p=0.3257).

### Differential effects of CSD

As we observed before (Pittaras et al., 2013; 2016), 3 groups of preferences emerged during the MGT. For the control group, we observed that only safe mice chose the advantageous choices more at the end of the MGT (safe: day1 *vs.* day 5, t=−4.923, p=0.007; average: day1 *vs.* day 5, t=−0.626, p=0.53; risky: day1 *vs.* day 5, t=−2.084, p=0.12, **Figure 3A**). However, the 3 groups differed from each other during the 2 last days of the MGT (day 1: H=2.732, p=0.25; day 2: H=4.482, p=0.10; day 3: H=5.140, p=0.07; day 4: H=11.371, p=0.003; day 5: H=11.690, p=0.002).

**Figure 2:**
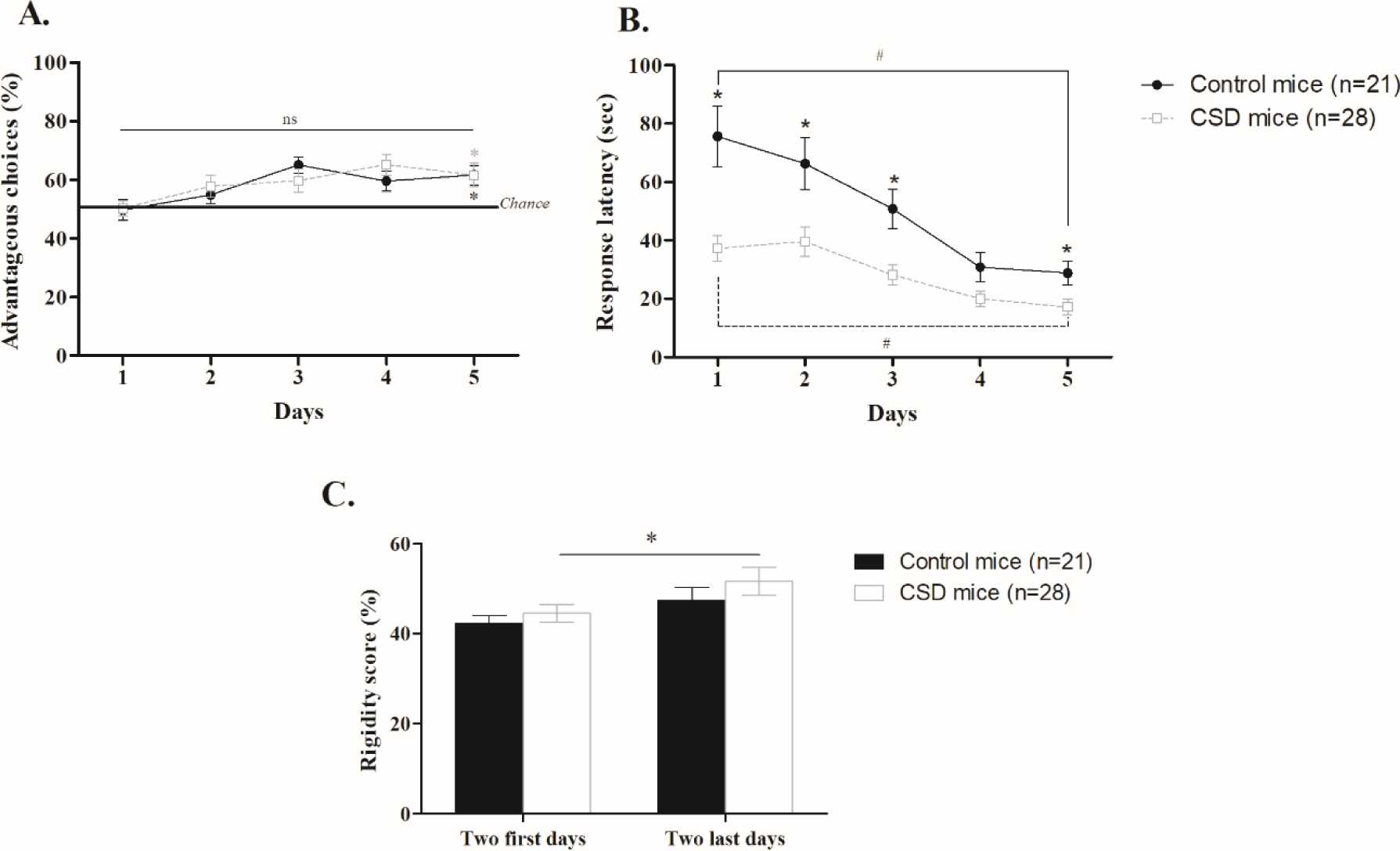
**A. Progression of mice preference during the Mouse Gambling Task (MGT)** for control (n=21, full black line) and Chronic Sleep debt mice (CSD, n=28, grey dotted line). Control and CSD mice choose more advantageous choices at the end of the task than at the beginning (*, p<0.05) but they never differed from each other (ns, p>0.5). **B. Progression of mice response latency during the MGT** for control (n=21, full black line) and CSD mice (n=28, grey dotted line). Control and CSD mice were quicker at the end of the task than at the beginning (#, p<0.05) and CSD mice were quicker than control mice for days 1, 2, 3 and 5 (*, p<0.05). **C. Rigidity scores** are reflected by the percentage of chosen arms during the 2 first days and the 2 last days of the MGT for control (black) and CSD (white) mice. Only CSD mice showed an increase in their rigidity at the end of the task compared to the beginning (*, p<0.05).

**Figure 3:**
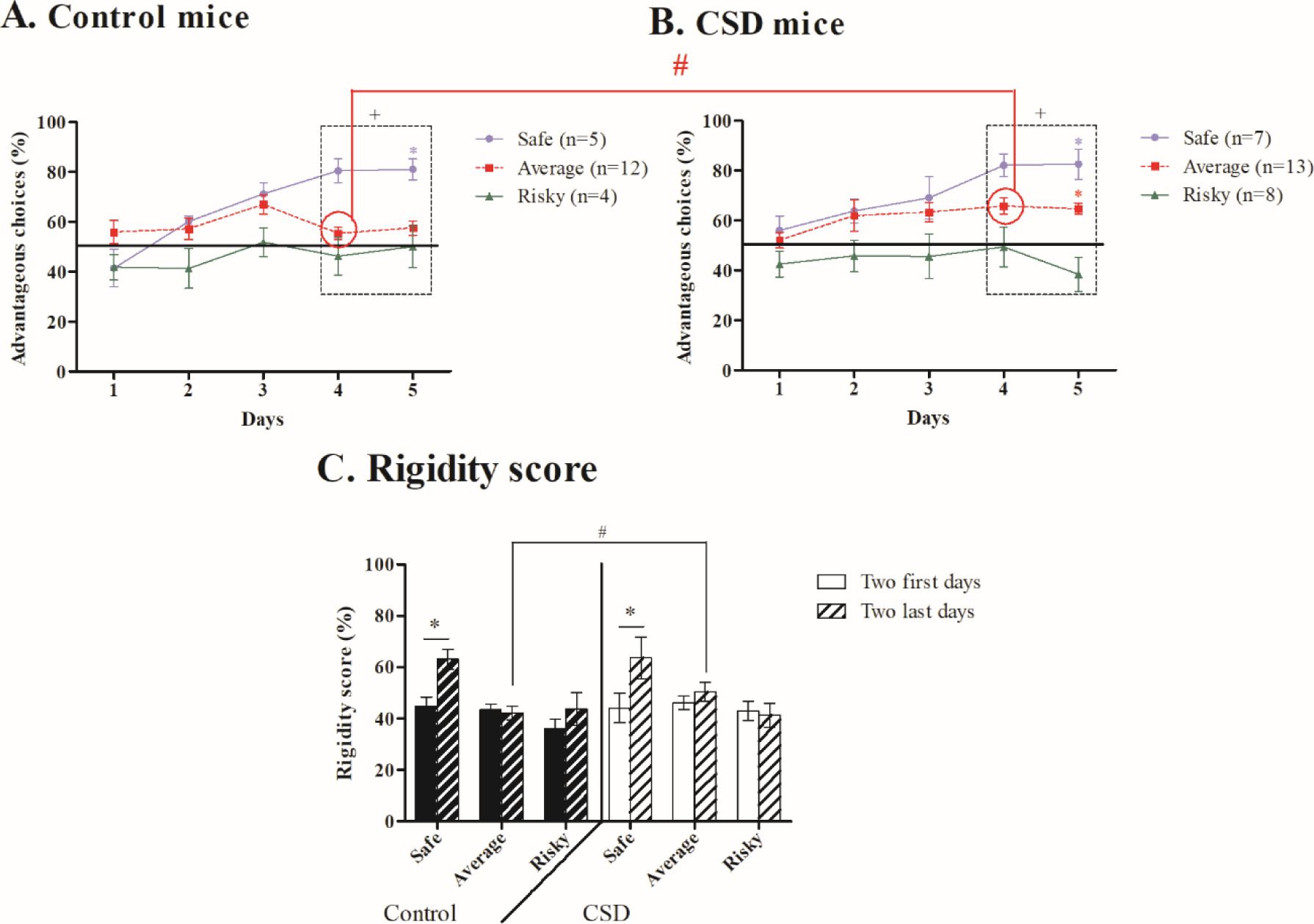
**A. Progression of mice preference during the Mouse Gambling Task (MGT)** for safe (n=5, blue circle line), average (n=12, dotted red line) and risky control mice (n=4, green triangle line). Only safe control mice chose advantageous choices more at the end of the task than at the beginning (*, p<0.05). The 3 groups differed from each other during days 4 and 5 (+, p<0.05). **B. Progression of mice preference during the Mouse Gambling Task (MGT)** for safe (n=7, blue circle line), average (n=13, dotted red line) and risky (n=8, green triangle line) Chronic Sleep Debt (CSD) mice. Safe and average CSD mice chose advantageous options more at the end of the MGT compared to the beginning (*, p<0.05). The 3 groups differed from each other during days 4 and 5 (+, p<0.05). **C. Rigidity scores** are reflected by the percentage of chosen arms during the 2 first days (filled) and the 2 last days (dotted) of the MGT for control (black) and CSD (white) mice. Only safe mice showed an increase in their rigidity at the end of the task compared to the beginning (*, p<0.05). Average control mice were less rigid than average CSD mice (#, p<0.05).

These inter-individual differences were also observed with the CSD mice. As shown in **Figure 3B**, safe and average CSD mice chose more advantageous choices at the end of the MGT compared to the beginning (safe: day1 *vs.* day 5, t=−3.624, p=0.01; average: day1 *vs.* day 5, t=−2.134, p=0.03; risky: day1 *vs.* day 5, t=0.425, p=0.68). These three groups differed from each other during the two last days of the MGT (day 1: H=3.974, p=0.13; day 2: H=5.004, p=0.08; day 3: H=4.619, p=0.09; day 4: H=12.764, p=0.0012; day 5: H=14.867, p=0.0006).

Interestingly, average control and CSD mice differed from each other during day 4 (t=−2.576, p=0.016).

Regarding their rigidity, only the safe mice became more rigid at the end of the MGT (safe control: t=−3.463, p=0.025; average control: t=0.346, p=0.73; risky control: t=−1.740, p=0.18; safe CSD: t=−2.028, p=0.04; average CSD: t=−1.511, p=0.15; risky CSD: t=0.325, p=0.75; **Figure 3C**). We also observed that average control mice were less rigid than average CSD mice at the end of the MGT (t=−2.182, p=0.03).

## Discussion

Our results showed that when mice completed the Mouse Gambling Task (MGT) under chronic sleep debt (CSD), they were able, as a group, to show a long term advantageous decision-making strategies. We observed before that an acute sleep debt (ASD) tended to prevent mice, as a group, to select long term advantageous decision-making strategies (Pittaras et al., 2018). This observation was true only when the ASD was applied between the exploration phase (day 1 and 2) and the exploitation phase (day 4 and 5), when mice just began to understand which option is better. Here, mice were under sleep debt during all the phases of the MGT and even before the task. Thus, we could have guessed that they would not be able to show advantageous long term decision-making strategies.

Our results could be explained by several hypotheses. First, we can hypothesize that the CSD has no impact on decision-making processes because 4 hours of sleep per day is not enough to alter decision-making strategies. It has been shown in humans that CSD (4 hours of sleep per night during 7 days) lead to the same attention and memory deficits as 2 nights of ASD (Van Dongen et al, 2003). We also know that 6 hours of sleep debt for 4 days impairs spatial memory and suppresses hippocampal neurogenesis in rats (Hairston et al., 2005). As the MGT requires spatial memory consolidation, we should observe a decrease of mice’s MGT performances under sleep debt. Therefore, it is unlikely that 4 hours of sleep are enough to preserve decision-making in mice, but this still needs to be tested.

A more likely second hypothesis could be that mice are able to adapt themselves to sleep debt and set up a compensatory mechanism during the week of CSD. It has been shown in humans that a 5 hours CSD for 7 days leads to attention deficits at the beginning and a then a performance stabilization (Belenky et al., 2003). Moreover, after 3 days of CSD, rat’s performance during the rat PVT increases to reach the same level of performance than animals not under CSD (Deurveilher et al., 2015). Deurveilher et al. proposed that this performance recovery could be explained by habituation of rPVT, an increase in micro-sleep quantity or an increase in slow waves during sleep periods. Particularly, the decrease of slow waves leads to a decrease of adenosine that has been shown to have deleterious effects during the rPVT (Deurveilher et al., 2015). Therefore, we can hypothesize that a compensatory mechanism occurred during the week of CSD leading to no alteration of decision-making in the MGT. However, more experimentation needs to be done to test this hypothesis.

Also, when mice are under CSD, their response latency decreases and their rigidity increases during the MGT. We have shown before that an ASD decreases response latency only after its application and increases rigidity at the end of the MGT (Pittaras et al., 2018). We have also shown that this observation could be linked with an increase in impulsivity and monoaminergic brain modification (Pittaras et al., 2016). We can thus hypothesize that ASD and CSD both increase impulsivity and rigidity via monoaminergic communication. Measuring the monoaminergic level in the different brain areas involved in decision-making after one week of sleep debt would allow us to test this hypothesis.

As we observed before (Pittaras et al., 2013; 2016; 2018) a Gaussian repartition of mice performances appears during the MGT. Interestingly, we observed this repartition for the CSD and the control mice. Moreover, the 3 decision-making profiles and their flexibilities were the same for the CSD and the control mice. The only differences we observed were that, under CSD, average control mice were not more rigid at the end of the task and chose less advantageous options than average CSD mice on the fourth day, even if they preferred advantageous options at the end of the MGT. We can thus hypothesize that the cylinder was more stressful for average control mice which lead to slightly different behavior, but not to decision-making deficits.

In conclusion, we have shown here that a CSD does not interfere with decision-making strategies likely because mice get use to CSD and set a compensatory mechanism to maximize the beneficial effects of sleep during the hours when sleep is possible. Because we observed before that an ASD of 23 hours at a specific time has deleterious effects on decision-making processes, we can hypotheses that it is more deleterious for decision-making in mice to lose one entire night than to sleep less during 1 week. It has been shown in human that the duration of sleep debt and cognitive deficits are linked. Indeed, 8 hours of sleep deprivation has more deleterious effects on memory than 8 hours of sleep debt applied for two days (4 hours on bed by night during two days, Drake et al., 2001). Therefore, our results confirm this observation. Even though we showed no effect of CSD on decision-making, we also showed that ASD and CSD have the same effect on mice as on humans during decision-making processes. This observation reinforces the fact that the rodent model is suitable to study the effect of sleep debt on cognitive deficit and, more interestingly, that the Mouse Gambling Task is a very suitable task to study such effects.

